# Post exam item analysis: Implication for intervention

**DOI:** 10.1101/510081

## Abstract

Post exam item analysis enables teachers to reduce biases on student achievement assessments and improve their way of instruction. Difficulty indices, discrimination power and distracter efficiencies were commonly investigated in item analysis. This research was intended to investigate the difficulty and discrimination indices, distracters efficiency, whole test reliability and construct defects in summative test for freshman common course at Gondar CTE. In this study, 176 exam papers were analyzed in terms of difficulty index, point bi-serial correlation and distracter efficiencies. Internal consistency reliability and construct defects such as meaningless stems, punctuation errors and inconsistencies in option formats were also investigated. Results revealed that the summative test as a whole has moderate difficulty level (0.56 ± 0.20) and good distracter efficiency (85.71% ± 29%). However, the exam was poor in terms of discrimination power (0.16 ± 0.28) and internal consistency reliability (KR-20 = 0.58). Only one item has good discrimination power and one more item excellent in its discrimination. About 41.9% of the items were either too easy or too difficult. Inconsistency in option formats or inappropriate options, punctuation errors and meaningless stems were also observed. Thus, future test development interventions should give due emphasis on item reliability, discrimination coefficient and item construct defects.

## 1. Background

Education quality in Ethiopia seemed to be compromised by the rapid expansion of higher education institutions in the country (4). According to Arega Yirdaw (2016), problems in the teaching-learning process were amongst the key factors in determining education quality in private higher institutions in Ethiopia. Within the teaching-learning processes, effective assessment tools have to be delivered to measure the desired learning outcomes.

It is advisable to use appropriate instruments for assessing students at higher institutions (5). The rational for employing effective assessment tool is that assessment of students’ achievement is an integral part of the teaching - learning process (2). Assessments should track each student’s performance in a given course. With this in mind, instructors at colleges and universities must be aware of the quality and reliability of their exams in a given course. Otherwise, the final results may lead to a biased evaluation and certification (5). Instructors usually receive little or no training on quality of assessment tools. Trainings usually focus on large-scale test administration and standardized test score interpretation but not on strategies to construct test or item-writing rules (2). The quality and reliability of assessments can be improved by delivering trainings on post exam item analysis and item writing rules (17).

Item analysis involves collecting, summarizing and using information from students’ responses for assessing the quality of test items (21). It allows teachers to identify too difficult or too easy items, items that do not discriminate high and low able students or items that have implausible distracters (2, 3). By analyzing items, teachers/instructors can remove too easy/difficult items, improve distracters’ efficiency and avoid non-discriminating items from the pool of future test banks. It will also help teachers/instructors to examine misconceptions or contents difficult to understand for students and adjust the way they teach (2).

According to the reports in Ethiopia, there was a serious problem in quality of education (4, 19). Student’s achievement grading system in Ethiopia is carried out by administering teacher made classroom tests and national examinations (20). It is believed that assessing students’ performance solely on objective items at school and national level in the country might have contributed negatively on education quality (20). Therefore, objective test items need to met psychometric standards in order to measure the out comes as per the course objectives. Researchers suggested that objective examination results can be analyzed to improve the validity and reliability of assessments (17). Therefore, the objective of this research was to analyze the post examination results of a summative exam in basic natural science course at Gondar CTE. Based on the results, areas for intervention in future test development were recommended.

## 2. Methods

### 2.1 Research Design

The validity and reliability of a summative test in a freshman common course entitled as ‘basic natural science-I’ was assessed using descriptive analytical method. Of the two approaches of item analysis, classical test theory (CTT) and item response theory (IRT), CTT was employed due to its simplicity and lack of software applications for IRT. The psychometric parameters considered in this study were difficulty indices, point bi-serial correlations, internal consistency reliability and construct face validity, and distracter efficiency.

### 2.2 Study population

All regular first year diploma students at Gondar CTE during 2017/18 academic year were taken as the study population.

### 2.3 Sample size and sampling technique

Assuming homogeneity of the population, a total of 176 (33.5%) students (Total = 525) were selected using stratified random sampling. A stratified sampling technique was employed to include representative samples from each department. The sample exam papers were collected from science instructors within the department, Natural Science, Gondar CTE. Demographic data of the representative samples were collected from registrar office in the college.

### 2.4 Instrument and scoring

The summative test administered during 2018 academic year in the course ‘basic natural science-I’ was used as the research instrument. The first reason why this course was selected for the study was that the summative test was developed by instructors with Biology, Chemistry and Physics educational backgrounds. Therefore, the findings would be applicable to the department, Natural Science. The other reason was that it was a common course given to a large population in the college as a compulsory common course to all new modality streams. Furthermore, the course was a pre-requisite for most other courses within the stream, integrated natural science. Therefore, it would be better if an effective assessment tool was prepared by the department. The summative test used in this study contained 31 objective items. There were 21 multiple choices, 7 true/false items and 3 matching questions. All the 31 items were considered for analysis. For item analysis, correct responses were coded as 1 and 0 for wrong responses. The maximum mark possible to score was 31 and minimum zero, with no negative marking.

### 2.5 Construct defect (Face validity)

The exam paper was checked for the following construct defects.

- Typing and punctuation errors

- Inappropriate/incomplete stems

- Inappropriate options formats/alternatives format for MCQs.

### 2.6 Internal consistency reliability

The internal consistency reliability of the summative test in basic natural science-I course was investigated to determine the overall reliability of the test. The two most commonly used measure of reliability were Cronbach alfa (*α*) and Kuder-Richardson method (KR-20). KR-20 is used to measure the reliability of tests in a dichotomous item (17). Therefore, in this study, KR-20 was used to estimate the test reliability. The objective test items were scored dichotomously as right or wrong (17). Every correct response was coded as 1 and wrong responses as 0. The acceptable value for test reliability in many literatures was *α* ≥ 0.7. A KR-20 value of 0.7 or greater was considered as reliable in this study.

### 2.7 Item difficulty index (*p*)

The item difficulty index is an appropriate choice for achievement tests when the items were scored dichotomously. It can be calculated for true-false, multiple choice and matching items. In this study, difficulty index for every item was determined by dividing the number of respondents who answered the item correctly by the total number of students taking the test. Simply, *p* was computed using Microsoft excel 2007 based on the formula given below and average difficulty level determined.

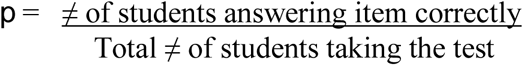

The value of *p* ranges from 0 - 1; the higher the value, the easier the item and vice versa. The recommended range of difficulty level is between 0.3 - 0.7 (1, 6). Items having *p*-values below 0.3 and above 0.7 are considered too difficult and too easy, respectively (1).

| Item difficulty index ($p$) | Item evaluation |
| --- | --- |
| $p > 0.7$ | Too easy |
| $p = 0.3 - 0.7$ | Acceptable |
| $p < 0.30$ | Too difficult |

Source: Instructional Assessment Resources (IAR 2011) in (1)

### 2.8 Discrimination coefficient (*r*)

The item discrimination index is a value of how well a question is able to differentiate between students who are high performing and those who are not (17). It can be calculated either by extreme group method or point bi-serial correlation coefficient (*r*) or other methods. The extreme group method considers only 54% of the respondents (top 27% and bottom 27%). On the other hand, the point bi-serial correlation coefficients take into account all respondents. Besides, it also indicates the relationship between a particular item on a test with the total test score (12, 17). For this reason, the point bi-serial correlation coefficient was used in this study. The point bi-serial correlation coefficient was computed using SPSS version 20. Its value ranges between −1 and 1; a higher value indicates a powerful discrimination power of the item. The test items in this study were classified based on the standard depicted in the table below.

| $r$ value | Quality | Recommendations |
| --- | --- | --- |
| $\geq 0.4$ | Excellent | Retain |
| 0.3 – 0.39 | Good | Possibilities for improvement |
| 0.2 – 0.29 | Average | Need to check/review |
| 0.0 – 0.19 | Poor | Discard or review in depth |
| < - 0.01 | Worst | Definitely discard |

Adopted from (7)

### 2.9 Distracter efficiency (DE)

Distracters are classified as incorrect answers in a multiple-choice question. Student performance in an exam is very much influenced by the quality of the given distracters. Hence, it is necessary to determine the effectiveness of distracters in a given MCQ. Distracter effectiveness indicates the percentage of students choosing that option as an answer. It was calculated based on the number of non-functional distracters (NFDs) per item. An NFD was defined as an incorrect option in MCQ selected by less than 5% of students. The DE was considered to be 0%, 33.3%, 66.7% or 100% if an item had three, two, one or zero NFDs, respectively.

## 3 Data analysis

The Statistical Package for Social Science software version 20 (SPSS-20) and Microsoft Excel 2007 were used for storing and analyzing data. Descriptive statistics i.e. frequencies, demographic information, mean and standard deviations as well as point bi-serial correlations were determined using SPSS-20. Difficulty indices and percentages were computed using excel. Face validity was described qualitatively. Figures and tables were used to display results.

## 4 Results

### Demographic data

Table 1 below shows the demographic characteristics of students whose exam papers were analyzed. Forty two percent of the samples were females and males constitute fifty eight percent. More than half (58.5%) of the study participants were in the age group 20-25 years; 38.1% were under 20 and only 3.4% were 26 and above. The mean age was 20.26 ± 2.13.

**Table 1.** Age-sex demographic characteristics of respondents

| Sex | Frequency | Percent |
| --- | --- | --- |
| Male | 102 | 58.0 |
| Female | 74 | 42.0 |
| Total | 176 | 100.0 |
| Age in years | Frequency | Percent |
| Under 20 | 67 | 38.1 |
| 20 - 25 | 103 | 58.5 |
| Above 25 | 6 | 3.4 |
| Total | 176 | 100.0 |

### Test statistics

Results from Table 2 showed that students’ score ranged from 5 to 27 (out of 31) with mean 17.23 ± 3.85. There was no statistically significant mean difference between males and females (p = 0.311, df =174). The histogram in figure 1 revealed that the total score was approximately normally distributed in both sexes (Fig 1).

**Table 2:**
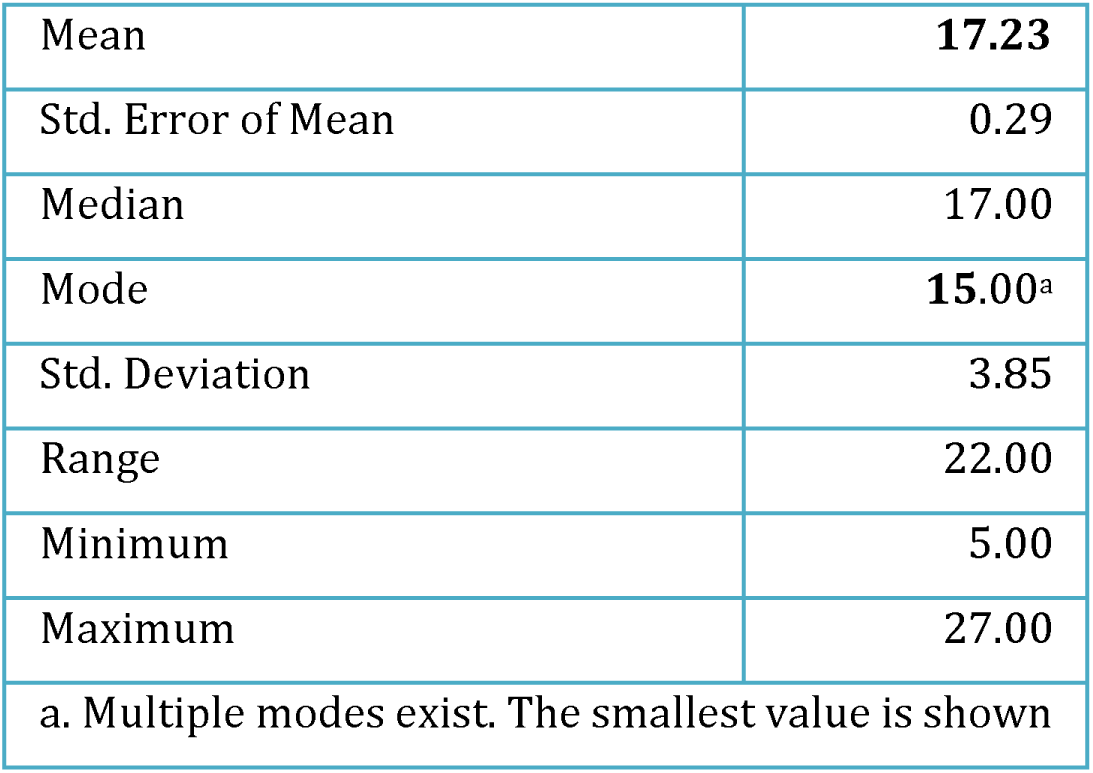
Item descriptive statistics

|  |  |
| --- | --- |
| Mean | **17.23** |
| Std. Error of Mean | 0.29 |
| Median | 17.00 |
| Mode | **15.00<sup>a</sup>** |
| Std. Deviation | 3.85 |
| Range | 22.00 |
| Minimum | 5.00 |
| Maximum | 27.00 |
| a. Multiple modes exist. The smallest value is shown |  |

**Figure 1.**
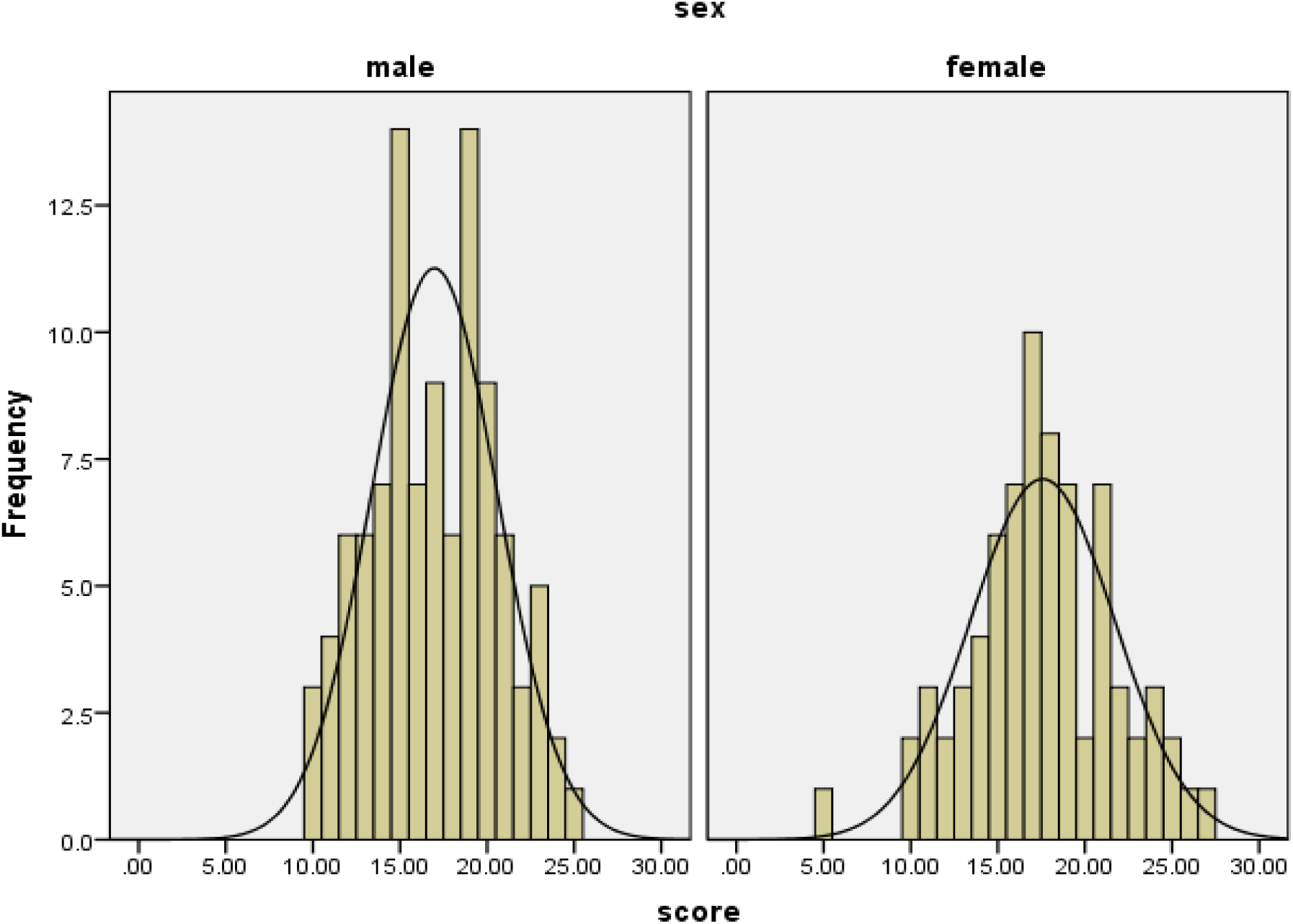
Distribution of raw test score in males and females

### Construct defects (Face validity)

Results from face validity revealed the following findings.

- Punctuation errors (full stop & question mark missing) in question.5, 6, 7, 11 and 15.

- Inconsistent option formats (option format changed from question 26 to 31).

- Inappropriate stems (meaningless or incomplete) were observed in question 12, 13, 14, 27 and 28.

- Inappropriate options/alternatives (all of the above, A and B) in question 12, 13, 15, 22, 27 and 28.

- No negatively phrased stems *(not* or *except).*

- Absolute term (most) in question number 4.

## Internal consistency reliability

In this study, the internal consistency reliability was used to evaluate the performance of the test as a whole. The computed KR-20 value of the test was 0.58 which is less than the recommended range in many literatures (≥ 0.7).

### Difficulty index

Appendix A shows the distribution of difficulty indices (*p*) for each item. One item (q.19) has the highest *p*-value (0.82) and q.18 has the lowest (0.15). Eighteen items (58.1%) have moderate difficulty level (*p*-value between 0.3 - 0.7). Twelve items (38.7%) have excellent difficulty levels (*p*-values between 0.4 - 0.6). Thirteen items (41.9%) lie outside the moderate difficulty range i.e. three items were too difficult (*p* < 0.3) and ten items (32.3%) too easy (*p* > 0.7). The mean difficulty index was 56% that is *p* = 0.56, SD. 0.20. A summary of difficulty index was illustrated in Table 3.

**Table 3:**
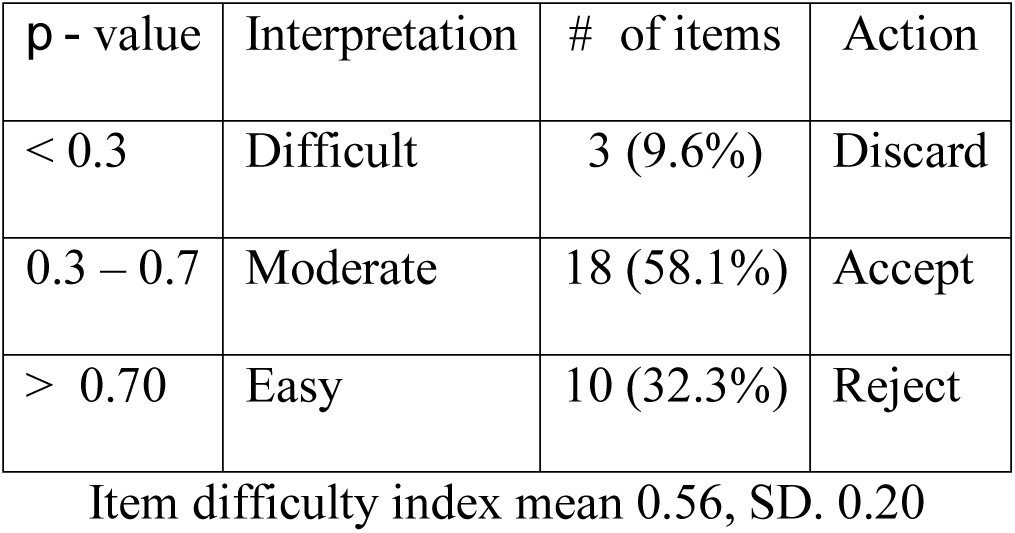
Difficulty index summary

| $p$ - value | Interpretation | # of items | Action |
| --- | --- | --- | --- |
| $< 0.3$ | Difficult | 3 (9.6%) | Discard |
| 0.3 – 0.7 | Moderate | 18 (58.1%) | Accept |
| $> 0.70$ | Easy | 10 (32.3%) | Reject |

Item difficulty index mean 0.56, SD. 0.20

### Discrimination coefficients

Appendix B shows the result of the point bi-serial correlation coefficient for each item. Three items (**q.2, q.18** and **q.20**) have negative discrimination power (*r*-worst). Only a single item (**q.31**) has excellent discrimination power (*r* > 0.4). Seventeen items (54.8%) were categorized as poor (*r* < 2.0) and nine items (29%) as average (*r* = 2.0 - 2.9) (Table 4). Question number **7** is an ideal item in terms of difficulty level (*p* = 0.54, Appendix A), but good in terms of discrimination (*r* = 0.39, Appendix B). The mean discrimination power is **0.16** (SD. 0.28). In many literatures, the acceptable mean *r* value is ≥ 0.4.

**Table 4:** Distribution of items in terms of level of discrimination

| Point bi-serial Correlation ($r$) | No. of Items | % | Action |
| --- | --- | --- | --- |
| Excellent ($r \geq 0.40$) | 1 | 3.23% | Retain |
| Good ($r = 0.30 - 0.39$) | 1 | 3.23% | Possibilities for improvement |
| Average ($r = 0.20-0.29$) | 9 | 29.03% | Usually need and subject to improvement |
| Poor ($r = 0 - 0.19$) | 17 | 54.84 | Discard or review in depth |
| Worst ($r < 0$) | 3 | 9.67% | Definitely discard |

Item discrimination coefficients mean 0.16, SD, 0.28.

Table 5 below displays the combination of the two indices i.e. item difficulty and discrimination.According to this table, two items (**q.7** and **31**) have moderate p-values (*p* = 0.3 - 0.7) and good discrimination (*r* ≥ 0.3). However, there was no any single item that could be labeled as excellent in both difficulty and discrimination indices (*p* = 0.4 - 0.6 and *r ≥* 0.4). Easy items (*p* > 0.7) such as **q.3, q.5, q.10** and **q.24** have poor discrimination (*r* < 0.2). Furthermore, items with *p* < 0.3 (difficult) such as q.12, q.14 and q.18 have very low discriminating power (*r* < 0.2). The difficulty level for q.2 was ideal (*p* = 0.51) but its discrimination power was worst (*r* = −0.004). Ideal questions with *p*-values from 0.4 to 0.6 (q.6, q.16, q.21, q.23 and q.26) have poor *r-*values (< 0.2). There was no statistically significant correlation between difficulty index and discrimination coefficient (Pearson correlation = 0.201, Sig. (2-tailed) p = 0.279).

**Table 5:**
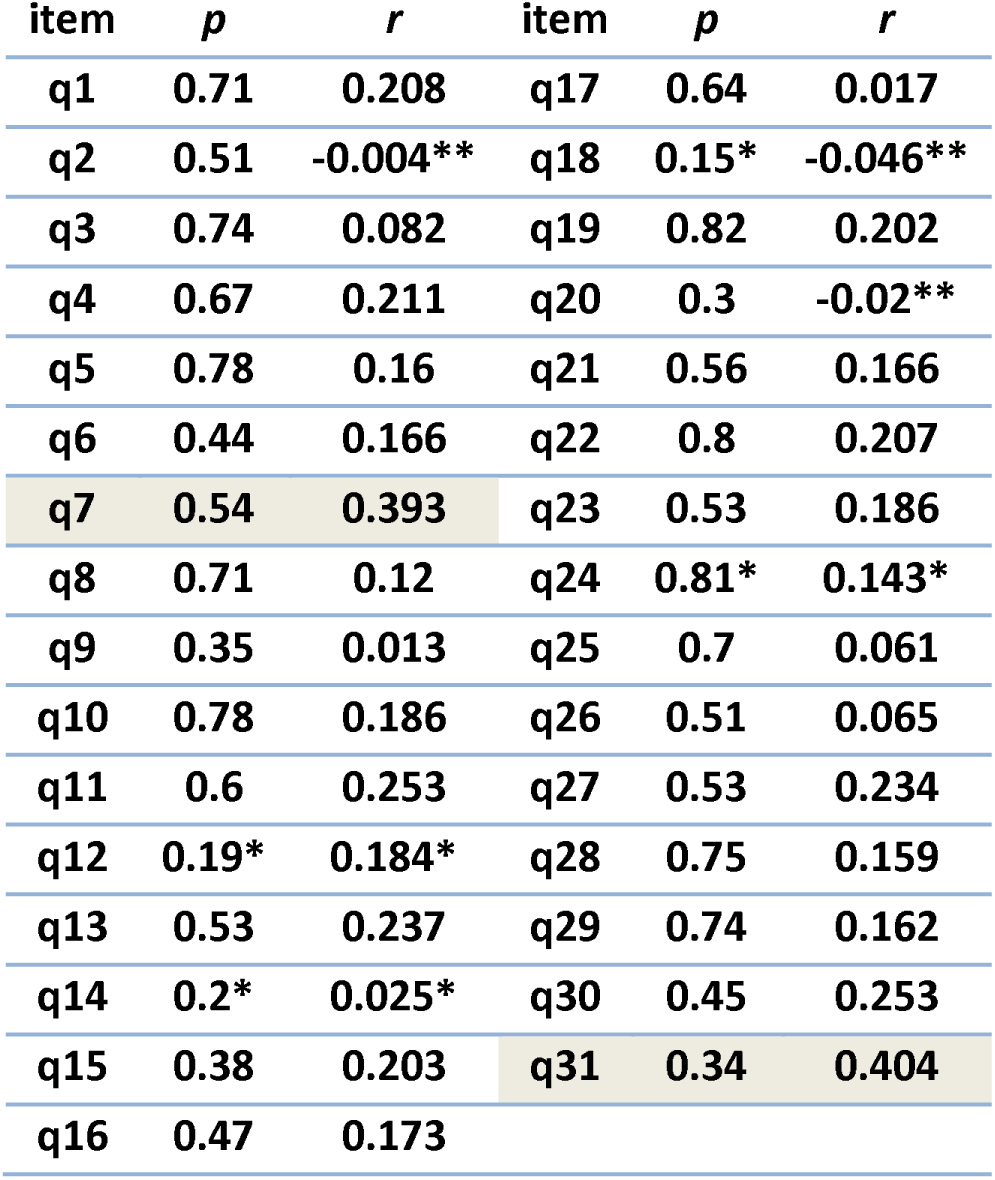
Combination of item difficulty (*p*) and discrimination (*r*) indices

| item | $p$ | $r$ | item | $p$ | $r$ |
| --- | --- | --- | --- | --- | --- |
| q1 | 0.71 | 0.208 | q17 | 0.64 | 0.017 |
| q2 | 0.51 | -0.004** | q18 | 0.15* | -0.046** |
| q3 | 0.74 | 0.082 | q19 | 0.82 | 0.202 |
| q4 | 0.67 | 0.211 | q20 | 0.3 | -0.02** |
| q5 | 0.78 | 0.16 | q21 | 0.56 | 0.166 |
| q6 | 0.44 | 0.166 | q22 | 0.8 | 0.207 |
| q7 | 0.54 | 0.393 | q23 | 0.53 | 0.186 |
| q8 | 0.71 | 0.12 | q24 | 0.81* | 0.143* |
| q9 | 0.35 | 0.013 | q25 | 0.7 | 0.061 |
| q10 | 0.78 | 0.186 | q26 | 0.51 | 0.065 |
| q11 | 0.6 | 0.253 | q27 | 0.53 | 0.234 |
| q12 | 0.19* | 0.184* | q28 | 0.75 | 0.159 |
| q13 | 0.53 | 0.237 | q29 | 0.74 | 0.162 |
| q14 | 0.2* | 0.025* | q30 | 0.45 | 0.253 |
| q15 | 0.38 | 0.203 | q31 | 0.34 | 0.404 |
| q16 | 0.47 | 0.173 |  |  |  |

Fig.2 shows the graphical representation of difficulty index and discrimination power. The scatter plot allows identification of appropriate (valid and reliable) questions at the center of the graph. Moreover the representation is useful to notice immediately questions that are too easy or too difficult. According to fig 2, *r* increases up to a point where *p* approaches to 0.4, then after it declines.

**Figure 2:**
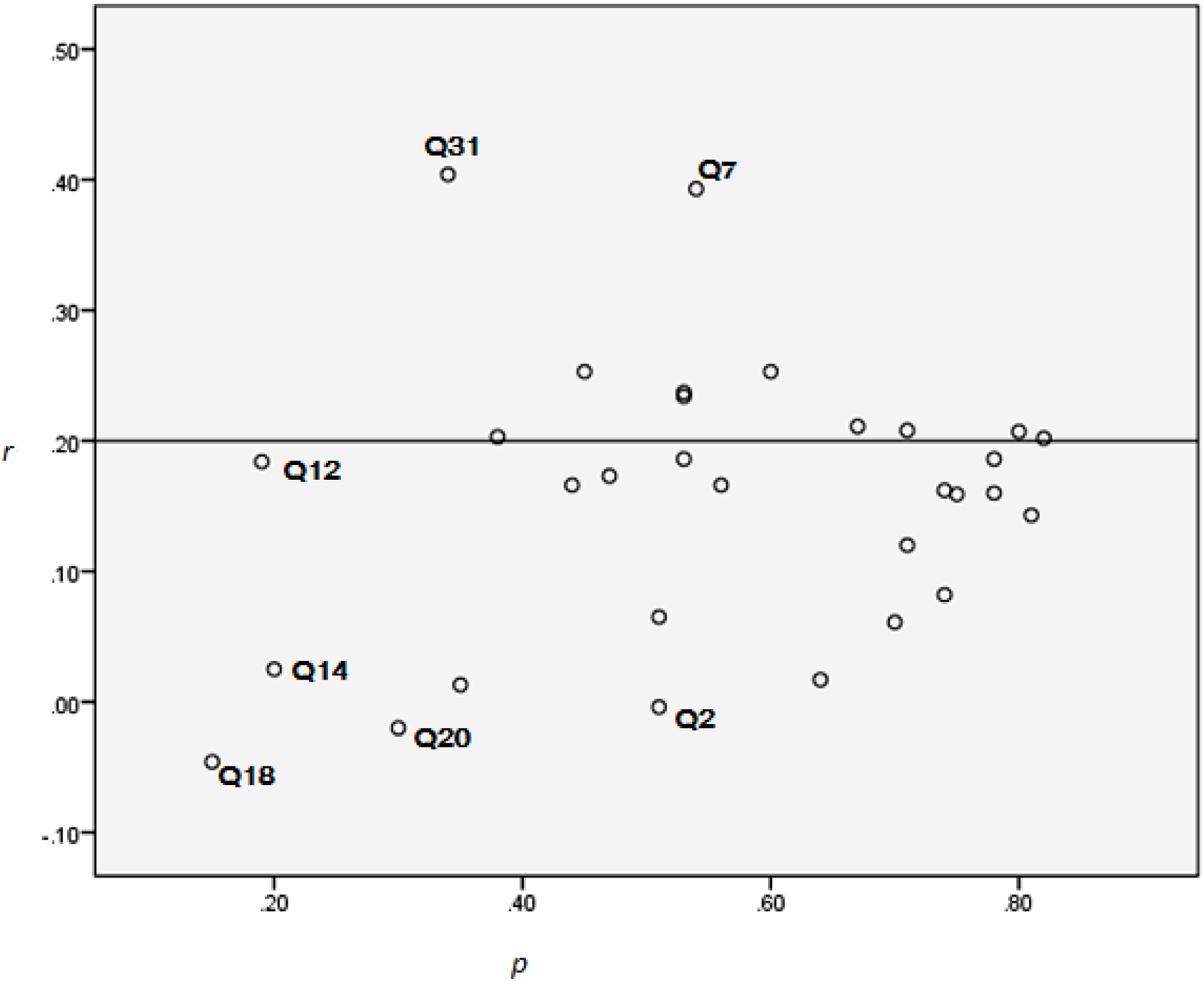
Scatter plot of difficulty and discrimination indices

### Distracter efficiency

The distracter analysis shows that six items (28.57%) (**q.12, q.13, q.19 q.20, q.24** and **q.28**) have nine NFDs, with a choice frequency of < **5**%. All other items do not have any NFDs (Appendix C). In addition, four items (**q.12, q.14, q.18** and **q.20**) have six distracters selected by more students than the (key) correct answer. There was no item with three NFDs. But three items (**q.12, q.13 and q.19**) have 2 NFDs and the next three items (**q.19, q.20** and **q.28**) have 1 NFDs (Table 6). The overall mean of DE was 85.71% with minimum 33.3% and maximum 100%.

**Table 6:** Distracter analysis (DE) summary

|  |  |
| --- | --- |
| Number of Items | 21 |
| Total Distracters | 63 |
| Functional distracters | 54 (85.71%) |
| Non functional distracters (NFDs) | 9 (14.29%) |
| Items with 3 NFDs (DE=0%) | 0 |
| Items with 2 NFDs (DE=33.3%) | 3 |
| Items with 1 NFD (DE=66.7%) | 3 |
| Items with 0 NFD (DE=100%) | 15 |
| Items with over distracters | 4 |
| Overall mean DE | 85.71% ± 24.89% |

Difficult items such as **q.12, q.14** and **q.18** have DE between 66.7% and 100%. Similar result was recorded for easy questions such as **q.18, q.22** and **q.24**. Some items with poor or good *p-*values have similar **DE** values (Table 7). Only a single item (**q.31**) satisfies all the three parameters of ideal questions (*p* > 0.3, *r* > 0.4 and DE = 100%, Table 8).

**Table 7:** Comparison of item difficulty with distracter efficiency for MCQsTable 8: Comparison of p, *r* and DE

| Item | *p* | DE (%) |
| --- | --- | --- |
| q11 | 0.6 | 100 |
| q12 | 0.19* | 66.7 |
| q13 | 0.53 | 33.3 |
| q14 | 0.2* | 100 |
| q15 | 0.38 | 100 |
| q16 | 0.47 | 100 |
| q17 | 0.64 | 100 |
| q18 | 0.15* | 100 |
| q19 | 0.82 | 66.7 |
| q20 | 0.3 | 100 |
| q21 | 0.56 | 100 |
| q22 | 0.8 | 100 |
| q23 | 0.53 | 100 |
| q24 | 0.81* | 100 |
| q25 | 0.7 | 100 |
| q26 | 0.51 | 100 |
| q27 | 0.53 | 100 |
| q28 | 0.75 | 66.7 |
| q29 | 0.74 | 100 |
| q30 | 0.45 | 100 |
| q31 | 0.34 | 100 |

**Table 8:** Comparison of *p, r* and DE

| item | $p$ | $d$ | DE |
| --- | --- | --- | --- |
| q11 | 0.6 | 0.253 | 100 |
| q12 | 0.19* | 0.184* | 66.7 |
| q13 | 0.53 | 0.237 | 33.3 |
| q14 | 0.2* | 0.025* | 100 |
| q15 | 0.38 | 0.203 | 100 |
| q16 | 0.47 | 0.173 | 100 |
| q17 | 0.64 | 0.017 | 100 |
| q18 | 0.15* | -0.046** | 100 |
| q19 | 0.82 | 0.202 | 66.7 |
| q20 | 0.3 | -0.02** | 100 |
| q21 | 0.56 | 0.166 | 100 |
| q22 | 0.8 | 0.207 | 100 |
| q23 | 0.53 | 0.186 | 100 |
| q24 | 0.81* | 0.143* | 100 |
| q25 | 0.7 | 0.061 | 100 |
| q26 | 0.51 | 0.065 | 100 |
| q27 | 0.53 | 0.234 | 100 |
| q28 | 0.75 | 0.159 | 66.7 |
| q29 | 0.74 | 0.162 | 100 |
| q30 | 0.45 | 0.253 | 100 |
| q31 | 0.34 | 0.404 | 100 |

## Discussion

By analyzing summative assessments, it is possible to modify future test development techniques or modify classroom instructions. With this intention the current study was conducted on post exam item analysis based on psychometric standards. Based on the findings areas for interventions were indicated.

The internal reliability calculated in this summative test was 0.58. This value is a beat less than the expected range in most standardized assessments (*α* ≥ 0.7). According to (8), a Cronbach alpha of 0.71 was obtained in a standardized Italian case study. Reliability analysis could be categorized as: excellent if *α* > 0.9, very good if between 0.8 - 0.9 and good if between 0.6 - 0.7 (1). If the reliability value lies within 0.5 - 0.6, revision is required. It will be questionable if reliability falls below 0.5 (1). Based on the result in this study, the summative test administered requires revision because its KR-20 (0.58) value is less than 0.7. This might also imply that college educators need to validate their assessment tools through item analysis. According to Fraenkel and Wallen in (12), one should attempt to generate KR-20 reliability of 0.70 and above to acquire reliable test instruments.

According to Table 3, 58.1% (18) of the items in the summative test have average difficulty (*p* = 0.3 - 0.7). An item is considered to be good item if its *p*-value lies in the moderate range (17). In this study, a little higher than half of the exam items have moderate difficulty. It is important to include more questions with average difficulty even though the mean difficulty level (0.56) is acceptable. A similar finding was reported in many other literatures (1, 6 and 9).

Questions which are too easy or too difficult for a student contribute little information regarding student’s ability (17). Data in this study showed that 32.3% of the test items were too easy (recommended – 10%-20%) and 9.6% were too difficult (recommended-20%). Though it is advisable to include easy and difficult items in a given test (10), it would be better if the recommended limits were met. Hence in this exam paper, there were more easy items and fewer difficult items. A difficult item could mean either the topic is difficult for students to grasp (10, 11) or not taught well (10) or mis-keyed (12) or poor preparation of students.

According to (12), the discriminatory power of individual items can be computed either by discrimination index, biserial correlation coefficient, point biserial coefficient or phi coefficient. In this study, the discrimination power of every item was calculated by using point-biserial coefficient. The point-bi-serial coefficient result (Table 4) showed that only one item was considered as ‘Excellent” (*r* > 0.4) and another one reasonably good (*r* = 0.393). All other items in this summative test need revision or subjected to improvement (*r* < 0.3). Similar study was reported in (3) that there was no a single item having discrimination index greater than or equal to 0.30. Contrary to this study, 46.67% of items have good to excellent discrimination power (*r* ≥ 0.3) (15).

Large and positive values are required for the point bi-serial correlation as it indicates that students who get an item right tend to obtain high scores on the overall test and vise-versa (8). An item with negative and/or low discriminating power needed to be considered in subsequent test development phases. In this study, three items have negative discrimination. This could be due to the fact that low ability students guessed the item right and high ability students suspicious of any clue less successful to guess (16). Items with negative discrimination decrease the validity of the test and should be removed from the collection of questions (10, 12, 13, 14 and 15).

Difficulty and discrimination indices are often reciprocally related. However, this may not always be true. Questions having high *p*-value discriminate poorly; conversely, questions with low *p*-value may discriminate well (17). This variation could be as a result of students who make a guess when selecting the correct responses (12). Data (Table 5) showed that guessing has occurred in this study. According to (1), moderately difficult items should have the maximal discriminative ability. The findings of this study contradict with (1). This may reflect that some extent of guessing occurred during test administration.

Distracter analysis was conducted to determine the relative usefulness of distracters in each item. Seventeen (81%) items have no NFDs (DE = 100%), three items (**q.12, q.19** and **q.28**) have 1 NFDs (DE = 66.7%) and one item (**q.13**) has 2 NFDs (DE = 33.3%, Table 6). There is no item with 3 NFDs (DE = 0). On the other hand, seven distracters (11%) (12 - A, 14 - A, B, 18- B, C and 20-A) were selected by more students than the correct answer. This may indicate that the items were confusing (12). An overall DE mean of 92.1% (considered as ideal/acceptable) was obtained in this study. Similar finding was reported by (10) in an internal microbiology examination in India.

Non-functional distracters make an item easy and reduce its discrimination (10). Question number 31 (Table 8) has moderate difficulty and excellent discrimination power. Probably this could be due to absence of NFDs. However, this doesn’t work for other test items probably due to random guessing or some flaws in item writing (10).

### 5.1 Conclusion and Recommendations

Post exam item analysis is a simple but effective method to assess the validity and reliability of a test. It detects specific technical flaws and provides information for further test improvement. An item with average difficulty (*p* = 0.3 – 0.7), high discrimination (*r* ≥ 0.4) and higher DE value (>70%) is considered as an ideal item. In this study, the summative test as a whole has moderate difficulty (mean = 0.56) and good distracter efficiency (mean = 85.71%). But it poorly discriminates between high and low achieving students (0.16). The test as a whole needs revision as its reliability was not reasonably good (KR-20=0.58). Some flaws in item writing were also observed.

According to Xu and Liu (2009) in (1), teachers’ knowledge in assessment and evaluation is not static but a dynamic and ongoing activity. Therefore, it is plausible to suggest that teachers or instructors should have some in-service seminars on test developments. Since most of the summative tests constructed within the college are objective types, item analysis is recommended for instructors at some points in their teaching life. It is also suggested that there might be a specific unit responsible for testing and the analysis of the items after exam administration.

## Acknowledgment

I would like to thank lecturers at Department of Natural Science, Gondar CTE, for providing exam papers for the study. I would like to extend my appreciation to Mr. Awoke Debebe for his assistance in data entry and critical review of the manuscript.

